# gyōza: a Snakemake workflow for modular analysis of deep-mutational scanning data

**DOI:** 10.1101/2025.02.19.639168

**Authors:** Romain Durand, Alicia Pageau, Christian R. Landry

## Abstract

Deep-mutational scanning (DMS) is a powerful technique that allows screening large libraries of mutants at high throughput. It has been used in many applications, including to estimate the fitness impact of all single mutants of entire proteins, to catalog drug resistance mutations and even to predict protein structures. Here, we present gyōza, a Snakemake-based workflow to analyze DMS data. gyōza requires little programming knowledge and comes with comprehensive documentation to help the user go from raw sequencing data to functional impact scores. Complete with quality control and an automatically generated HTML report, this new pipeline should facilitate the analysis of time-series DMS experiments. gyōza is freely available on GitHub (https://github.com/durr1602/gyoza).

**Article summary:** Deep-mutational scanning (DMS) refers to molecular biology methods used to generate many genetic variants and evaluate their effect on adaptive fitness. It is used both in fundamental and applied research, with implications in genomic medicine. Many data processing steps are needed to transform the output of DMS, high-throughput sequencing data, into scores that quantify the fitness effect of each variant. The analysis is tailored to the type of experimental design, which can vary a lot. To facilitate such analysis and make it accessible to people with limited knowledge in bioinformatics, we have developed gyōza, a free and easy-to-use program to analyze DMS data.

## 1. Introduction

Deep-mutational scanning (DMS) is a molecular biology technique that evaluates the functional impact of a large number of mutations introduced at a particular locus (Hietpas *et al*. 2011; Araya and Fowler 2011; Fowler and Fields 2014). Owing to its broad range of applications and its relatively low cost, DMS has gained in popularity (Wei and Li 2023). Once the library of mutants is constructed, it is expressed in cells and subjected to screening (for example by applying selective pressure, Figure 1).

**Figure 1.**
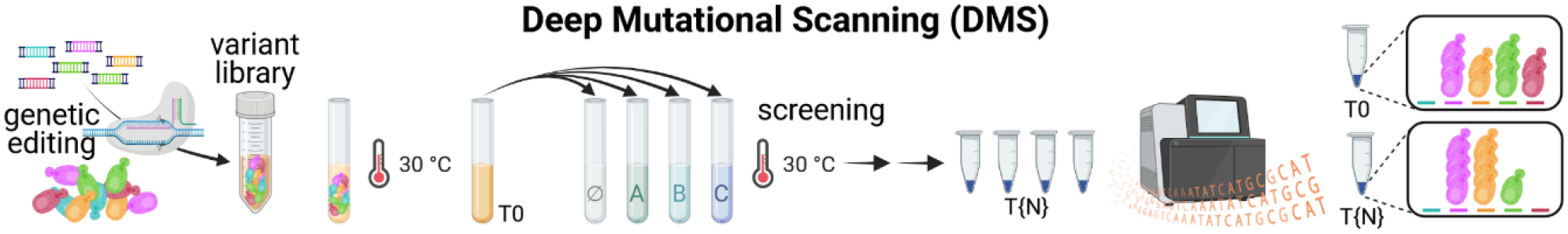
Overview of the typical experimental DMS workflow. Deep mutational scanning can be done many ways. A schematic example is represented here. First, variant libraries are generated for every locus of interest, for example, by genetic editing. Then, libraries are cultured to express the variants. Non-deleterious variants are sequenced from samples collected at this initial time point (T0), as well as after screening, usually in a control condition and one or several other conditions (e.g. A, B, C). Here, screening is done through successive rounds of selection, with sampling at each time point. Ultimately, the loci of interest are sequenced at high throughput. Figure created with BioRender.com, modified from Fig 1B of (Durand *et al*. 2024). **Alt text**. Schematic representation of a deep-mutational scanning experiment.

Comparing allele frequencies at any post-screening time point to the initial time point allows the estimation of mutational effects. Several tools have been developed to perform this task (Çubuk *et al*. 2025). Enrich2, one of the most popular DMS analysis tools, was developed early as a statistical framework to perform error estimation (Rubin *et al*. 2017). It has only recently been updated to be able to run on modern versions of Python. DiMSum is a more recent pipeline, written in R, which was also developed as a statistical framework to help users identify bottlenecks and other sources of error in their DMS dataset (Faure *et al*. 2020). Although DiMSum cannot handle multiple loci or multiple post-screening time points, it is well maintained and the report it generates is informative. Other recently published tools include Rosace (Rao *et al*. 2024), a Bayesian framework to analyze growth-based DMS data and popDMS (Hong *et al*. 2024), a DMS analysis method based on population genetics theory. Existing tools can be integrated into a single pipeline. For example, Dumpling is a Snakemake workflow that integrates both Rosace and Enrich2, allowing the use of either scoring method (Rao *et al*. 2024). For a more complete review of variant scoring software for DMS, we redirect the reader to a recently published article (Çubuk *et al*. 2025).

Despite the diversity of existing tools to analyze DMS data, there are still experimental designs (and platform restrictions in certain cases) that preclude their use. For example, at the time of writing, the Rosace environment can make it difficult to install and use on some systems, while popDMS uses a Gaussian prior distribution which is not suited for experiments involving strong selective pressure. This is typically the case for experiments involving proteins whose mutants confer antimicrobial resistance (Després *et al*. 2024; Chen *et al*. 2024; Bédard *et al*. 2024). In general, developing a DMS analysis pipeline that can suit anyone’s needs comes with a lot of challenges. Different experimental designs will lead to different sources of error, which means any statistical analysis is often project dependent. We believe that an interesting compromise is a pipeline with a simple core that produces estimates of mutational effects ready for custom statistical analyses. Additionally, it should be usable by anyone with limited bioinformatic knowledge. To that aim, we designed gyōza, a flexible and easy-to-use pipeline to analyze time-series DMS data. To make sure that gyōza is accessible, we implemented several quality checks to help the user prepare everything needed for a workflow run. This includes detailed warnings in case of formatting issues or if any file is missing. gyōza is a Snakemake workflow, which means it comes with all advantages related to Snakemake, such as standardized code organization, increased reproducibility and portability. Other Snakemake-related aspects have previously been detailed elsewhere (Mölder *et al*. 2021; Corut and Wallace 2023).

gyōza is easy to install and comes with a toy dataset to quickly check proper installation. For an actual run, the user needs to edit a short configuration text file, set up a few parameters and provide the necessary files depending on the experimental design. gyōza can handle several types of experimental designs including standard designs (mutations introduced at each position or at specific positions), barcoded designs and random mutagenesis data. Multiple loci can be analyzed separately in a single workflow run, regardless of the number of replicates or time points. An interactive HTML report is generated dynamically at the end of any run. This report will inform the user on how the sequencing data were converted, with plots and detailed captions for each step, optionally including an embedded interactive MultiQC report to explore the sequencing quality of all selected samples. Using this report, the user will know quickly if most of their sequencing reads passed all filters, or if they were lost at any of the early steps such as trimming or merging. If the user enables processing the read counts, these are converted into functional impact scores, reflecting mutational effects. If normalization with growth data is enabled, the calculated scores correspond to selection coefficients. The user can restrict the analysis to specific samples and/or choose which ones will feature in the report. gyōza can be run on a personal computer with limited computing resources or on typical high-performance computing (HPC) clusters, with almost no change in the setup. Most importantly, with comprehensive documentation and transparent source code, we hope to convince experimentalists to analyze their own DMS data.

## 2. Methods

### Setup

gyōza can be installed on Linux and Mac systems or WSL2 for Windows users, by following a few steps on the command line, including installing the necessary dependencies. Integrated package management with Snakemake currently relies on Conda (Anaconda 2016) (more details in the “Conda” section in the supplemental material), which means its installation is a prerequisite. To be able to install and use gyōza, we recommend the creation of a virtual environment, in which Snakemake, Snakedeploy and a few other dependencies will be installed. Snakedeploy is the official command line tool for deployment and maintenance of Snakemake workflows. Once the virtual environment is ready and activated, a single command line deploys gyōza on your system. A minimal file tree is automatically created and a message prompts the user to set up the configuration (see section below). This deployment is specific to a chosen version of gyōza, which increases reproducibility. The user can also initialize the created directory as a (public or private) git repository, in order to track changes, specifically in case of downstream analyses.

gyōza comes with a toy dataset that is automatically deployed, which means the installation can be quickly tested by running the same command as for an actual run (see “Local execution of the workflow” section). For a standard use, the toy dataset can be replaced with real data by adapting the configuration and providing the necessary input files.

### Configuration

The configuration file is a YAML file, which can be edited just as any text file, and is also validated against a provided YAML schema. This is particularly useful to ensure that no entry is missing and that all have the proper format. The most important config entry relates to the design of the DMS experiment. The user can choose between four options (at the time of writing):

- codon: for one or several mutated loci, the nucleotide sequences of all expected mutants based on a specified type of degenerate codon introduced at each position are automatically generated and annotated (currently supports single mutants with NNN or NNK at each position or corresponding pairwise combinations).
- provided: the nucleotide sequences of all expected mutants for each mutated locus are provided by the user.
- barcode: the dataframes of barcode-variant associations for each mutated locus are provided by the user.
- random: variants observed in the sequencing dataset are filtered based on a specified number of amino acid changes.

Other entries include:

- the path to the directory containing the project-specific files
- the project-specific sample attributes that feature as columns in the sample layout,
- the full path to the directory containing the sequencing reads (with an entry to specify if the reads are paired-end or single-end),
- parameters used for filtering and plotting purposes (expected read count per sample, T0 read count threshold to label variants, desired formats for exported graphs),
- options to enable/disable quality check of all raw sequencing reads using FASTQC and MultiQC, conversion of read counts into functional impact scores or normalization with the number of cellular generations.

### Project-specific files

The most important project-specific files correspond to demultiplexed sequencing data (FASTQ) and the sample layout. Sequencing data are expected to have been obtained with an Illumina or Aviti sequencer, can be paired-end or single-end and can optionally be compressed (.fastq.gz). Because it is often more convenient to keep sequencing datasets in a specific location, the user is asked to specify the full path to the directory (which may contain other files). For other project-specific files, they should be placed in a single folder. The user can either replace the files of the toy dataset in place or provide the full path to the prepared directory. For all designs, two files are mandatory: the sample layout and the codon table.

The sample layout is a CSV file containing the following mandatory columns:

- Sample_name: unique identifier for each sample,
- R1 and, optionally, R2: name of the demultiplexed FASTQ files containing the forward and reverse reads, respectively,
- N_forward and, optionally, N_reverse: 5’-3’ constant sequences on either side of the mutated locus,
- Mutated_seq: unique identifier for each mutated locus,
- Pos_start: starting position in the protein sequence for the mutated locus,
- Replicate: e.g. 1, 2 (more than 2 is handled but may lead to overplotting),
- Timepoint: T0 is mandatory, then one or more time points such as T1, T2, etc
- Analyze: “y”, “yes”, etc to select the sample for processing, meaning the data will be generated,
- Report: “y”, “yes”, etc to select the sample for processing and include it in the HTML report.

The “Mutated_seq” column is a convenient specificity of gyōza, compared to existing DMS analysis pipelines. It was introduced so that the user can analyze DMS data from different loci in a single workflow run. This is particularly useful for experiments in which a gene has been divided into several overlapping fragments (also called “tiles”). In such an experimental design, one would perform the DMS experiment on each fragment separately, then normalize the scores using the mutations in the overlaps. Because this additional normalization step is required only in certain rare cases (e.g. TileSeq), it is currently not included in the default workflow. Instead, we offer the possibility of labeling samples with any combination of attributes (e.g. attributes related to the fragment, the genetic background, the screening condition, the sequencing run etc). These attributes should be entered as additional column names in the sample layout (and should be listed in the configuration file).

Other project-specific files may be required depending on the design. ‘Codon’ and ‘random’ designs require that the user provides a dataframe containing the wild-type nucleotide sequences of each mutated locus. The CSV file should contain the following columns:

- Mutated_seq: identifiers matching the ones in the sample layout,
- WT_seq: corresponding wild-type DNA sequence, assuming the first three bases constitute the first mutated codon

For ‘codon’ designs, a third “codon_mode” column is expected, in which the user should specify what type of degenerate codon should be introduced at each position to generate the expected mutants.

‘Provided’ and ‘barcode’ designs require instead that the user provides dataframes containing all nucleotide sequences of expected mutants. One CSV file per mutated locus is expected, with the following columns:

- Mutated_seq: a single value per file (out of those listed in the Mutated_seq column of the sample layout)
- WT_seq: corresponding WT DNA sequence (single value per file), assuming the first three bases constitute the first mutated codon

For ‘barcode’ designs, a third “barcode” column is mandatory to list the barcode sequences corresponding to the nucleotide sequence of each expected mutant. Similarly to sample attributes, barcode attributes can be added to label the scores with barcode-level information such as barcode index, confidence related to variant association, etc). This information will be preserved in the full dataframe of non-aggregated functional impact scores.

If the user indicates in the config that they wish to normalize with the number of cellular generations, a template CSV file is automatically generated with the following columns:

- one column per sample attribute,
- Replicate (same as in the sample layout),
- Timepoint (here, only post-screening time points),
- Nb_gen (initialized with “1” in the template)

When the workflow is run for the very first time (including if it is a dry run), an exception is raised and a message displays on the terminal to notify the user that the template file has been generated and needs to be edited. The user will then edit the Nb_gen column with the number of cellular generations between T0 and the indicated time point on the same row, e.g. at “T2”, write “8.52” in the Nb_gen column to indicate that 8.52 doublings were measured between T0 and T2 for this condition. An example of the procedure applied to the toy dataset is provided in Figure S3.

All input files (except the sequencing data) are validated against dedicated YAML schemas to make sure that all mandatory columns are present and that the values in each column respect the expected format. If any formatting issue arises, detailed warnings should guide the user for troubleshooting.

In summary, for every new project, the user should edit the configuration file (YAML) and provide the following project-specific files:

- the FASTQ files of the paired-end sequencing reads,
- the sample layout (CSV),
- the wild-type sequences (CSV) for ‘codon’ and ‘random’ designs,
- the expected sequences (CSV) for ‘provided’ and ‘barcode’ designs,
- the number of generations between time points (CSV, based on generated template) if normalization enabled,
- the codon table if different from the provided one (*Saccharomyces cerevisiae* TAXID 559292)

### Structure and deployment of Snakemake workflows

Before we describe how to execute the workflow, let us introduce a few important concepts related to the use of Snakemake. A Snakemake workflow connects different modules, known as “rules”, which process input files and optionally generate output files. gyōza follows a standardized folder structure, with rules defined in Snakemake files (.smk) located in the “rules” directory. Each Snakemake file is imported in the “Snakefile”, depending on the configuration (e.g. the trimming stage requires different sets of arguments if reads are paired). The Snakefile additionally defines the main rule (“rule all”), which specifies the ultimate target file(s). Snakemake essentially works backwards from the ultimate target file(s) to build the workflow, connecting modules so that the necessary input files for each step are generated by an upstream module. Rule definitions include input, output, a main statement related to the actions and optional arguments such as parameters, resources, log files, etc. In summary, whenever an execution is triggered from the root directory (in our case, “gyoza”), Snakemake searches for the Snakefile and builds the workflow, starting with the main rule. Snakedeploy conveniently deploys only what is necessary, i.e. the files that need to be edited by the user. By abstracting the source code from the user, we prevent the introduction of errors, minimize the number of files copied locally and generally simplify the user experience. Since gyōza still needs some level of access to the source code, Snakedeploy packages the workflow into a module. Users may notice the creation of a minimal Snakefile upon deployment, which is necessary to run a Snakemake workflow, but in this case should contain very little information. The deployed Snakefile essentially specifies from which version of gyōza the code should be accessed. This allows for beta-testing, facilitating troubleshooting in some cases, in collaboration with the maintainers of gyōza. On the development side, maintainers can also use Snakedeploy to automatically update dependencies.

### Local execution of the workflow

In order to quickly check the integrity of the workflow (for example to ensure that filenames for the FASTQ files have been properly entered in the sample layout), it is strongly recommended to perform a “dry run”. This is done with a simple command line. At this stage, the workflow is essentially “pre-built”, with the output on the terminal listing all rules that are about to run (based on the config, existing input and output files, etc), including details for each job (which files will be created, why, see Figure S1). Most validations are performed during this dry run to make sure formatting errors, or errors related to the absence of required files, are spotted before the jobs are even submitted.

Local execution is also done through a simple command line and will automatically re-include a dry run. A “profile”, specified as a YAML configuration file is used by default and should require no modification by the user. Upon first use of gyōza, dedicated Conda environments will be installed and activated on demand to make sure the jobs of each rule are run in environments with the appropriate packages (more details in the “Conda” section in the supplemental material).

Snakemake cannot know the expected content of output files and will therefore not execute the workflow if the target files, identified by their path and name, already exist. On the other hand, some intermediary files are conserved by default, which means that Snakemake will always run whatever is “missing” for the workflow to complete. For example, if the workflow is accidentally aborted halfway through, rerunning the same command line will automatically detect that some files already exist and move on to the ones that are either incomplete or that have not been generated yet.

By default, all locally available cores will be provided. The user can scale down by specifying the number of CPUs to use in parallel. Note that if you are on a shared node, or if the dataset is rather large, it is strongly recommended to submit the jobs to the SLURM scheduler by specifying to use the SLURM profile (see “Running the workflow on HPC clusters” section). We typically recommend running the workflow locally for very few samples, then scaling up by switching to the SLURM profile. An example of the file tree generated upon execution is shown in Figure S2.

### Workflow steps

The main steps of the workflow are shown in Figure 2.

**Figure 2.**
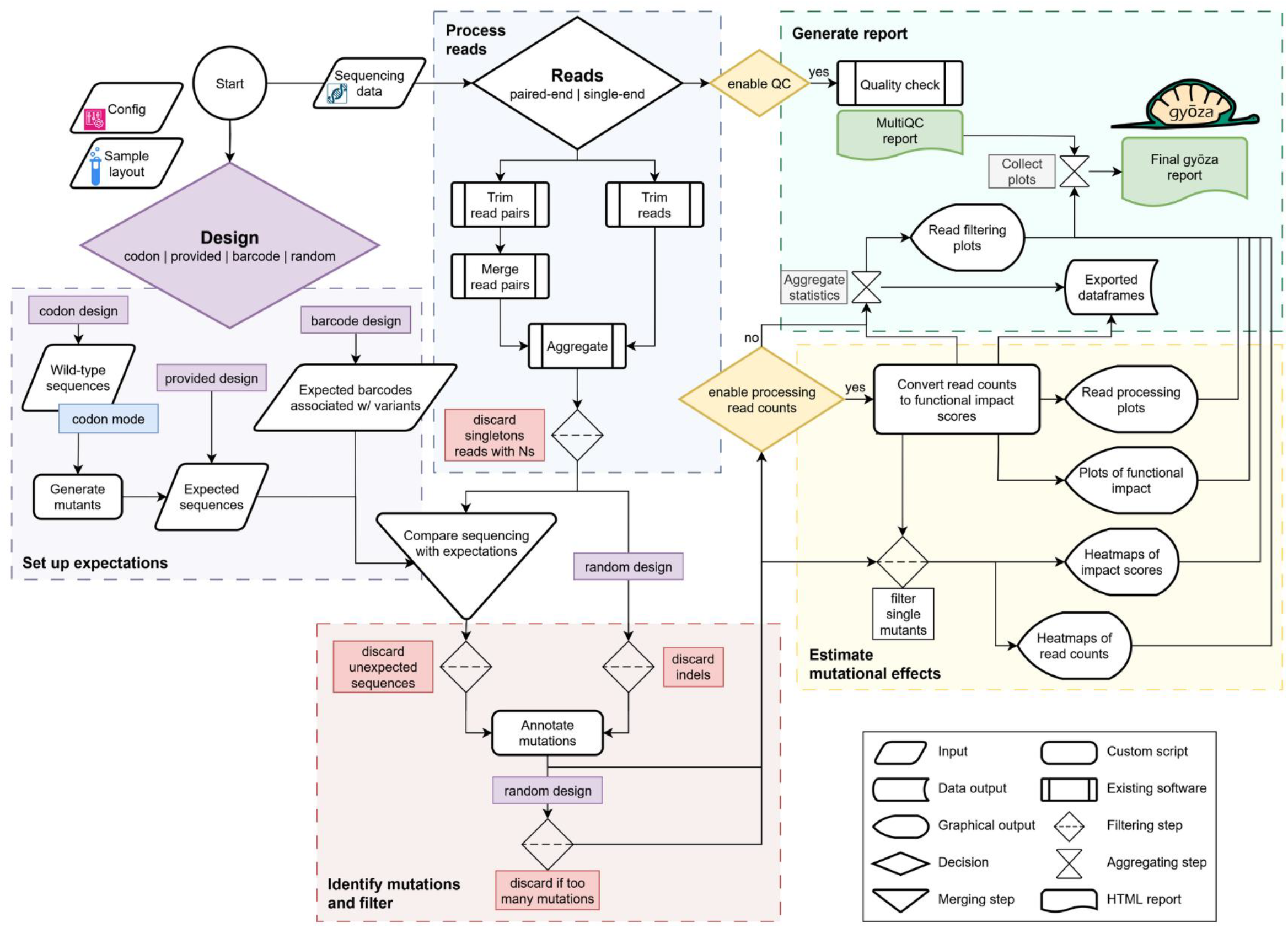
Overview of the gyōza workflow, depicted as a process flow diagram. The beginning of the workflow differs depending on the type of DMS experimental design specified in the config (highlighted in purple). For ‘codon’, ‘provided’ and ‘barcode’ designs, expected sequences are either generated or provided by the user (light purple box). For all designs, raw sequencing reads are processed using existing software. This includes an optional quality check with FASTQC, adapter trimming with Cutadapt, read merging using PANDAseq and aggregating reads for unique sequences using VSEARCH (light blue box). Sequences both “expected” and observed in the sequencing dataset are compared to the wild-type DNA sequence to identify mutations. For ‘barcode’ designs, the processed reads correspond to barcodes, while the expected sequences are barcodes previously associated with variants. Mutations are therefore identified by comparing the associated variants to the wild-type DNA sequence of the mutated locus (light red box). At this stage, the user can easily retrieve the dataframes of annotated read counts (one per sample). Several read statistics are collected at this stage and a visual representation is included in the final report (light green box). If the user has enabled processing read counts, a custom script converts them into functional impact scores using several normalization steps, taking into account sample depth, allele frequency at the initial time point, synonymous codons as a reference point and optionally the number of cellular generations (light yellow box). At the end of this step, two main dataframes are saved, the first with DNA-level or even barcode-level information and the second with protein-level information obtained by aggregating scores from high-confidence variants. A single HTML report is generated upon completion of the workflow, featuring plots with detailed captions, organized in sections, to estimate the overall quality of the dataset as well as get a preliminary overview of the results. Several steps, including steps related to export of intermediate dataframes, do not feature in this diagram to save space **Alt text**. Process flow diagram depicting the DMS analysis workflow implemented by gyoza.

#### Quality control

If enabled, raw FASTQ files are analyzed using FastQC (Andrews 2010) to evaluate the overall quality of sequencing data. A single interactive report is generated using MultiQC (Ewels *et al*. 2016) (and later on embedded in the gyōza report).

#### Getting read counts

Up to three steps are necessary to go from demultiplexed sequencing data to read counts: adapter trimming, read merging (for paired-end reads only) and counting the number of reads for each unique sequence. gyōza uses three packages to do that: Cutadapt (Martin 2011), PANDAseq (Masella *et al*. 2012) and VSEARCH (Rognes *et al*. 2016).

First, reads are trimmed with Cutadapt using specified constant sequences as adapters. This trimming stage is relatively flexible (5’ trimming, 3’ trimming, optional adapters, etc) to accommodate custom designs. By default, indels are forbidden and the maximum error rate is 0.15 (which means three substitutions in a 20-bp sequence would still match the adapter. The untrimmed sequences are discarded, but their number is retrieved by a downstream script (see “Gather read statistics” section). Trimmed forward and reverse reads are merged with PANDAseq. Among other technical details (which are transparent in the source code), we specify that the maximum overlap between forward and reverse reads from a pair be 625 bp. This is in line with PANDAseq recommendations in case of highly overlapping sequences, which is typically the case for Aviti sequencing reads. Finally, merged or trimmed single-end reads are collapsed into 100% identity clusters with VSEARCH (referred to as “dereplication” in the VSEARCH manual, with the function --fastx-uniques). In this final step, we discard all singletons (sequences only sequenced once), since they might result from sequencing errors. The FASTA output from VSEARCH is parsed separately for each sample using a custom script, which filters out sequences with uncalled bases and converts each FASTA into a CSV file with raw read counts.

#### Variant filtering

For all design options except ‘random’, a custom Python script simply uses the merge function from the pandas package to compare the observed sequences with the ones expected, essentially filtering out unexpected sequences, including sequences with incorrect length. The remaining sequences are then annotated with mutations inferred by comparing codons at matching positions with those of the wild-type DNA sequence. For ‘random’ designs, variants are directly annotated following read processing, then filtered based on the specified number of amino acid changes compared to the wild-type protein sequence. Variants that did not feature in the list of expected sequences and/or did not have the correct length, or had too many non-silent mutations, are all labeled “Unexpected” and are outputted to separate dataframes (one file per sample).

#### Gather read statistics

A custom Python script is used to parse all log files from upstream processing steps and output a single dataframe for all samples. This dataframe records how many reads passed all filters, or were discarded at each step (trimming, merging, aggregating, unexpected, contain Ns). To facilitate interpretation, this dataframe is plotted as a stacked bar plot, which ultimately features in the automatically generated HTML report (see “Exploring the results” section). The rationale is that the user can refer to this plot to quickly determine the quality of the dataset and resort to “manual” troubleshooting if need be. Some issues could be easier to troubleshoot than others: for example, if many reads are discarded at the trimming stage, the user might identify an error in the specified adapters. On the other hand, if a large proportion of reads are attributed to unexpected sequences, the user might need to manually inspect the content of discarded sequences.

#### Processing read counts

A custom Python script is used to process read counts, with the ultimate goal to convert them into functional impact scores. First, variants are labeled with a confidence score based on the number of reads at T0: 1, high confidence = above specified threshold in all replicates; 2, medium confidence = above threshold in at least one replicate; 3, low confidence = equal or below threshold in all replicates. The general rule to set a relevant threshold is to find the balance between:

- setting the threshold too low, which results in the inclusion of variants likely resulting from sequencing errors,
- setting the threshold too high, which diminishes the size of the dataset and may mask the full fitness range

In the very last steps of the workflow, gyōza saves a first dataframe with all functional scores and a second one including only average scores (and error) from high confidence variants across replicates.

The following steps are performed to convert read counts into functional impact scores:

- add 1 to all read counts,
- calculate read frequencies (normalize with sample depth),
- calculate log2 fold-change (frequency at a later time point replicate divided by frequency in the matching replicate at T0),
- (optional) normalize with the number of cellular generations between T0 and a later time point), in which case the ultimate score will correspond to a selection coefficient,
- subtract the median of normalized log2 fold-change values obtained for variants encoding the wild-type amino acid sequence, except for the wild-type nucleotide sequence

We purposefully exclude the wild-type nucleotide sequence from the last normalization step because it is often over-represented in libraries generated using oligonucleotide pools. Even with different experimental designs not relying on oligonucleotide pools, normalizing with a single reference may shift estimates of mutational effects. At this stage, non-aggregated scores are saved to CSV files (“all_scores.csv”) in the “results/df” directory. Each file contains the data from all samples sharing the same set of conditions (replicates and time points pooled together). Fitness values (average estimates of mutational effects) are obtained by calculating the median functional impact score for each amino acid sequence, from high-confidence nucleotide variants only. Finally, we calculate the median and error across replicates. The resulting dataframe is saved (“avg_scores.csv”). In this last table, for each output time point, three columns “fitness”, “lower_err” and “upper_err” correspond to the median, lower and upper error (from 2.5^th^ to 97.5^th^ percentile), respectively.

### Exploring the results

Throughout the workflow, plots are generated and stored as SVG files in the “graphs” directory, as well as in any format specified by the user in config. In order to facilitate interpretation of the analysis, a comprehensive HTML report is generated at the root of the “results” directory. This report will be generated regardless of the state of completion of the workflow, which means only completed steps will be included. It contains a main view with information related to the config and a minimal diagram of the workflow, as well as a side panel on the left with a search bar and a menu organized in sections (Figure 3a). Overall, the user has access to the following:

- custom message to describe how the specific dataset was meant to be analyzed (based on the config)
- source code for each rule with a list of input files, output files, parameters and packages with specified versions,
- runtime statistics,
- four main sections for the results:
  - Quality control: embeds the interactive MultiQC report on all FASTQ files (in case of display issues, the MultiQC report is also accessible separately),
  - Read filtering: stacked bar plot of filtered reads, distributions of read counts for unexpected variants in each sample and heatmaps of raw read counts for each sample,
  - Read processing: distribution of raw read count and frequency per variant, overlap across time points and replicates, distribution of allele frequencies at each time point,
  - Functional impact: distribution of functional impact scores, correlation between replicates, correlation between time points, functional impact over time and heatmaps of functional impact.

**Figure 3.**
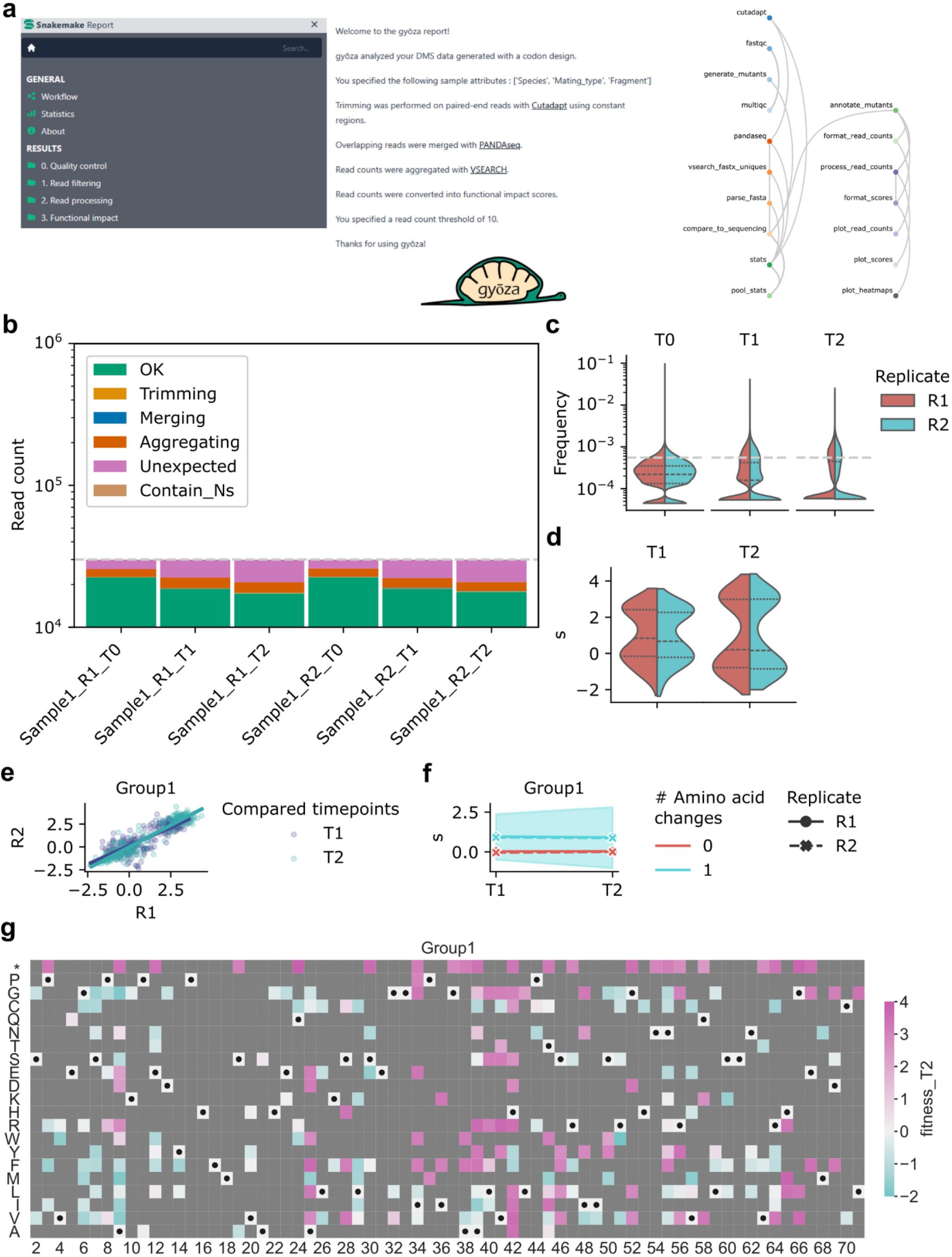
Examples of output generated by gyōza. a) Main view of the HTML report. Navigating the menu on the left will lead to several plots with detailed captions, including the examples in panels b-g. In the center, important information related to the config is displayed, as well as a simplified diagram of the workflow listing every rule that was run, with clickable icons to display the source code. b) Filtered reads. For each sample, read depths are plotted as stacked bars. The color key indicates the number of reads that passed all filters (“OK”), or the number of reads lost at the indicated step. c) Distribution of allele frequencies (read count per variant normalized by sample depth) at each time point. d) Distribution of functional impact scores at each post-screening time point. e) Correlation between replicates. f) Functional impact over time. Bands indicate the standard deviation for variants leading to the indicated number of amino acid changes compared to the wild-type protein sequence. g) Heatmap of functional impact scores at T2. The scores reflect the average impact from high-confidence variants only. The toy dataset was designed to have less reads than a typical dataset (for portability purposes), which means many variants do not meet the default read count threshold to be labeled with high confidence, ultimately leading to missing data on this specific heatmap. Plots have been slightly altered in Inkscape compared to the ones generated when running gyōza on the toy dataset, to maximize space efficiency. In general, the format of generated plots might vary depending on the number of samples and their properties (names, number of replicates, time points, etc). **Alt text**. Examples illustrating the report generated by the workflow, here with plots corresponding to the provided toy dataset.

Each plot comes with a detailed caption to help interpret the data. Examples are shown in Figure 3.

### Running the workflow on HPC clusters

If the user needs to scale up the workflow, gyōza supports the automatic submission of jobs to the SLURM scheduler. The user should first edit the SLURM profile located in the “profiles” directory, then run the appropriate command line to launch the workflow (specifying to use the SLURM profile instead of the default one). The SLURM profile is again a YAML file that can easily be modified by the user and constitutes a replacement for the bash files that are commonly used to submit jobs to SLURM. A user familiar with this procedure will find standard arguments in the sbatch statement, including a required email address. Make sure you keep only what is necessary and edit the arguments in the “default-resources” section. As their name indicates, these are only the default resources, in case some of them are not specified in the rule definitions. However, in order to prevent improper allocation and simplify user experience, gyōza defines rule-specific resources for demanding rules, which are dynamically requested based on the total input size and number of attempts. Priority will always be given to command line arguments over arguments specified in the profile. Other convenient options (implemented by default) include a naming scheme for every job that should make it easier for the user to find the proper log files if any error should occur. Log files related to SLURM jobs are located in the “logs/SLURM_out” directory and sorted by rule. They should contain the same output as what is usually displayed on the terminal and therefore may inform on errors related to Snakemake or SLURM. Errors related to existing software or custom scripts (not SLURM) feature in the single Snakemake log that is created for each run and whose path is indicated in one of the latest lines printed on the terminal.

By default, an email is sent to the specified email address every time a job fails (can be changed by modifying the “mail-type” argument in the profile). Conveniently, errors related to a lack of resources (job either reaches the time limit or is killed by an out-of-memory event) will be labeled accordingly in the subject of the email.

### 3. Results and discussion

gyōza comes with a toy dataset so that the user can test the installation. The same dataset was analyzed with DiMSum (Faure *et al*. 2020), although we selected only the first two time points (T0, T1), since DiMSum does not handle multiple post-screening time points. Overall, the setup is quite similar, except the preparation of input files for gyōza is intuitive, with validations and warnings providing helpful indications to set up the workflow. For a local execution (i.e. a small dataset), runtime is comparable once Conda environments for gyōza have been installed. However, scaling up with gyōza should prove fast both to set up and run. Internal tests with a 114 GB unpublished dataset (many conditions, including several time points, replicates and mutated loci) ran in 3.3 hours with a relatively small amount of computing resources (40 CPUs per task maximum at any time). This included FASTQC on 192 samples (384 FASTQ files), for a total of 2280 jobs.

Ultimately, most sequences (99.5 %) were picked up by both tools. Only 18 out of 3,867 sequences were discarded as singletons by one tool but not the other and vice versa (all with a read count of 2 instead of 1). Despite the differences in calculating the functional impact scores (DiMSum implements an error model), the values calculated by both tools correlate significantly (Figure 4). Unfortunately, attempts at testing the dumpling pipeline (which uses Rosace and Enrich2) were unsuccessful because of platform issues, which are being addressed at the time of writing.

**Figure 4.**
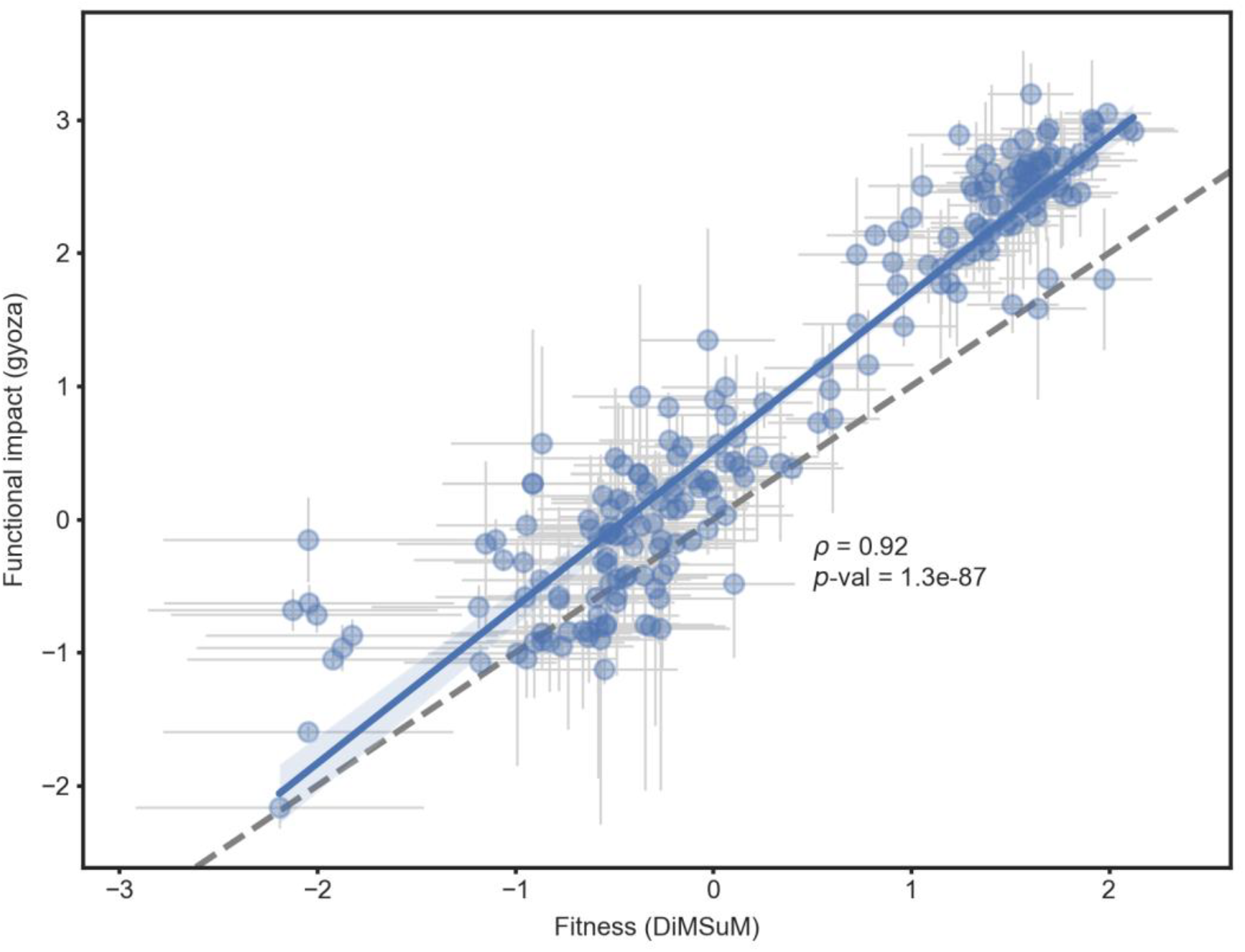
Correlation of functional impact scores calculated by gyōza v1.1.6 and DiMSum v1.3.2. Each data point corresponds to an amino acid sequence from the gyōza toy dataset. Functional scores (with 95 percentile interval as asymmetric error bars) calculated by gyōza are indicated on the y axis. “Fitness” and “sigma” (error) values calculated by DiMSum are shown on the x axis. Unique variants which were flagged as singletons (at T0 or T1) by either gyōza (n=1) or DiMSum (n=17) were excluded (in both cases, these 18 singletons were attributed a read count of 2 by the other tool). For both tools, scores were initially calculated for 3,849 unique nucleotide variants with a single mutated codon relative to the wildtype sequence. Average fitness values were calculated only from high confidence variants (i.e. variants from the toy dataset with a read count above 10 in all T0 replicates), resulting in 217 unique amino acid sequences. Correlation was estimated by calculating the Spearman’s rank correlation coefficient (ρ) and the corresponding *p*-value. Deviation from the diagonal can be explained by the use of different methods in normalizing the scores. **Alt text**. Graph comparing mutational effects, as estimated by the workflow and a similar existing tool.

In summary, gyōza provides a portable and intuitive way to quickly analyze time-series DMS data.

## 4. Data availability

gyōza is freely available (under MIT license) from GitHub (https://github.com/durr1602/gyoza). A separate repository contains everything that relates to tests reported here (https://github.com/Landrylab/Durand_et_al_2025_gyoza).

## 5. Acknowledgments

We thank Philippe C. Després for the initial optimization of PANDAseq and VSEARCH arguments as well as critical reading of the manuscript, Stéphane Larose for technical assistance with HPC and Victor Haffreingue for precious advice on the use of version control. We also thank members of the lab for insightful discussions and crucial feedback.

## 6. Funding

Canadian Institutes of Health Research (CIHR) (Foundation grant 387697, CRL)

Genome Canada and Genome Quebec (grant 6569, CRL)

Canada Research Chair in Cellular Systems and Synthetic Biology (CRL)

Natural Sciences and Engineering Research Council of Canada (NSERC) postdoctoral fellowship (RD)

## 8. Supplemental material

**Supplementary Figure 1.**
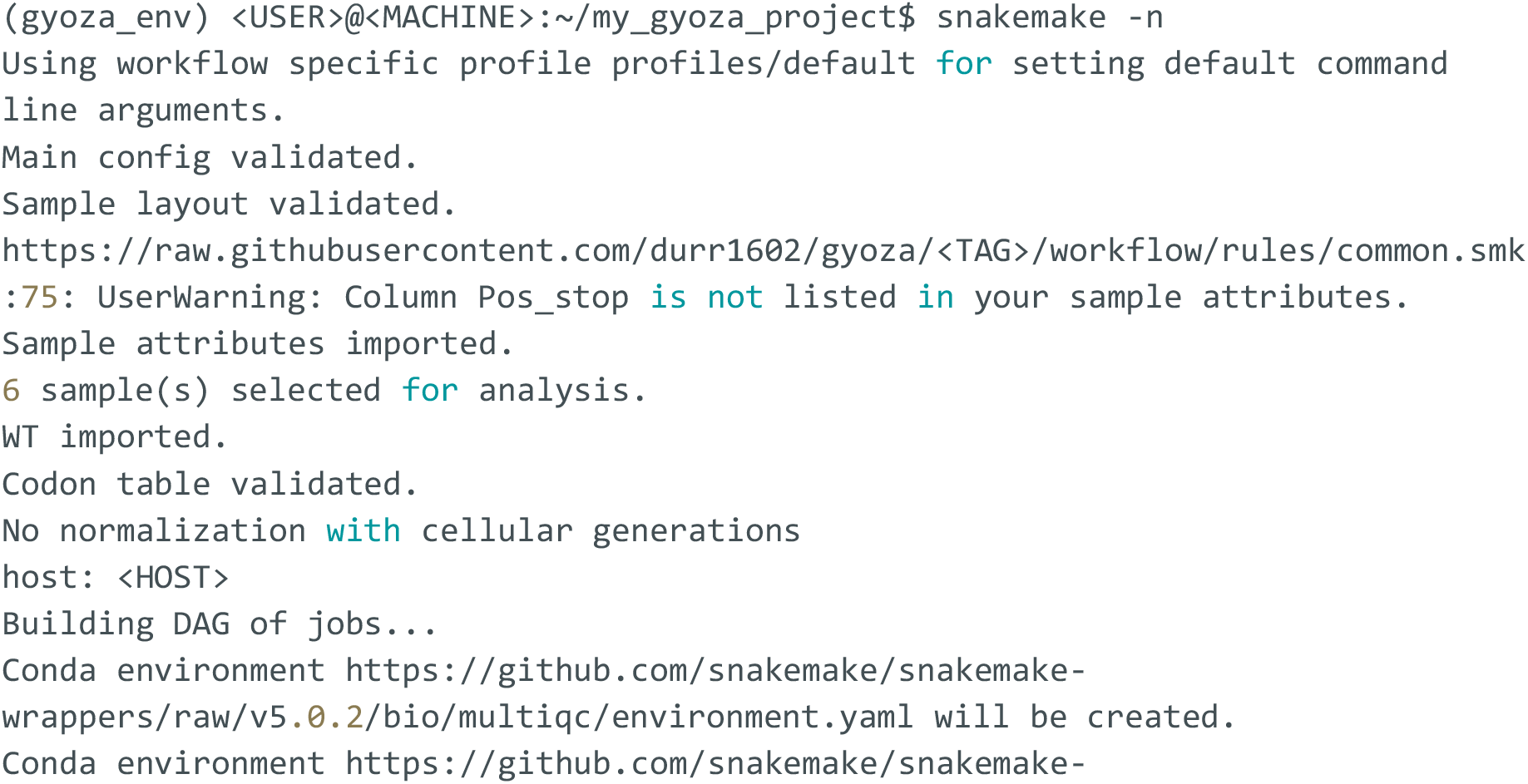
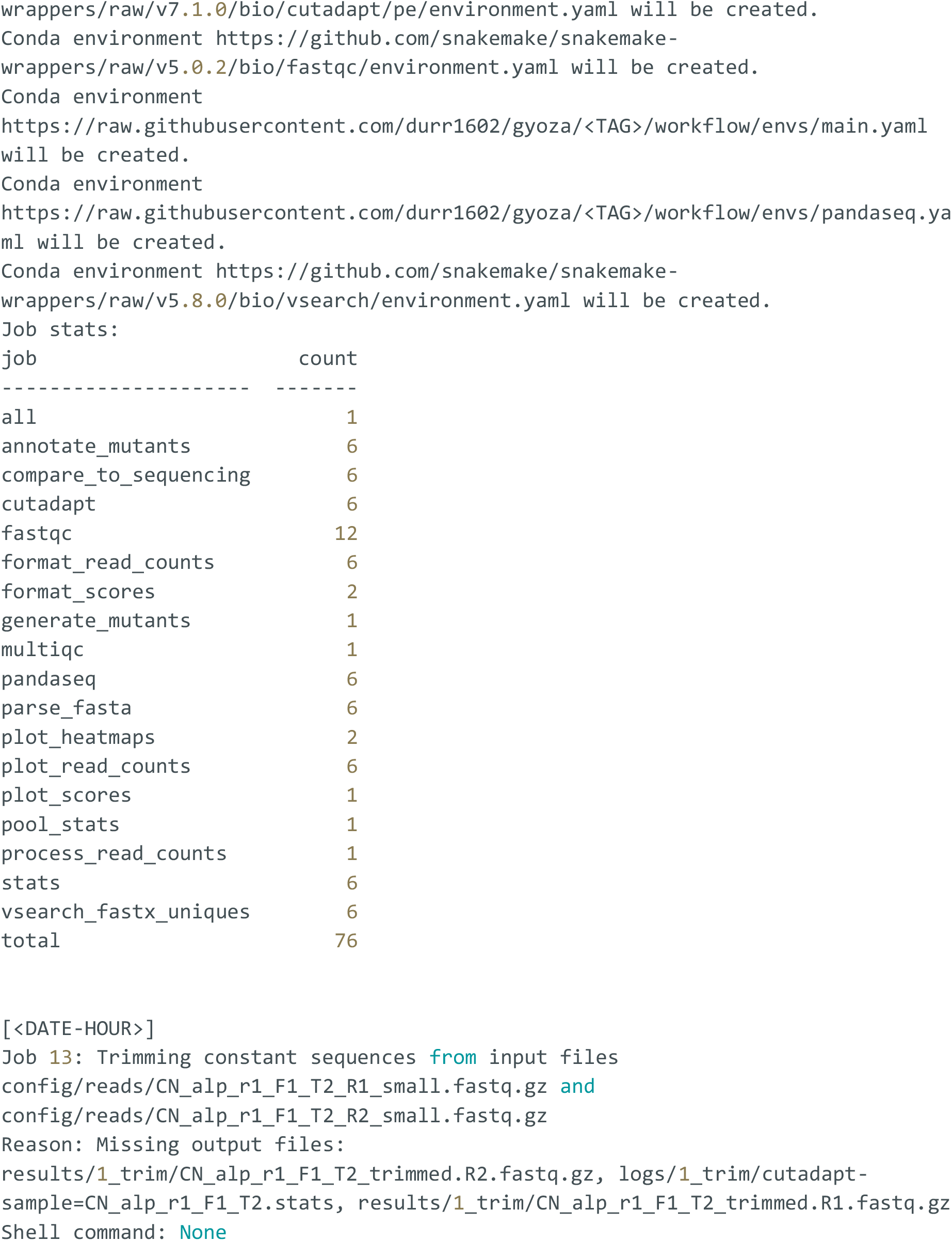
Terminal output for a dry run on the toy dataset. We show the beginning of the output printed to the terminal upon running the appropriate command line for a dry run. The very first line indicates which profile was used (either default for a local execution, or profiles/slurm if it was specified). A few lines correspond to validations of the config and the project-specific files. Finally, the workflow is built, the creation of rule-specific Conda environments is announced, as well as the corresponding rules (listed by alphabetical order), each with their job count. In the toy dataset, there are 6 samples selected for processing (as stated during the validations), which means there are 6 jobs for each rule processing samples separately. FASTQC runs on each FASTQ file separately, which means for paired-end reads, there should be twice as many jobs. For the rule “generate_mutants”, there is a single job, corresponding to the unique mutated locus in the toy dataset. Here, we don’t show the description of all 76 jobs, but we show the first one listed as an example (notice that jobs are not listed in order). A custom message describes what will be done during the corresponding job and why (missing output files, input files have changed, etc).

**Supplementary Figure 2.**
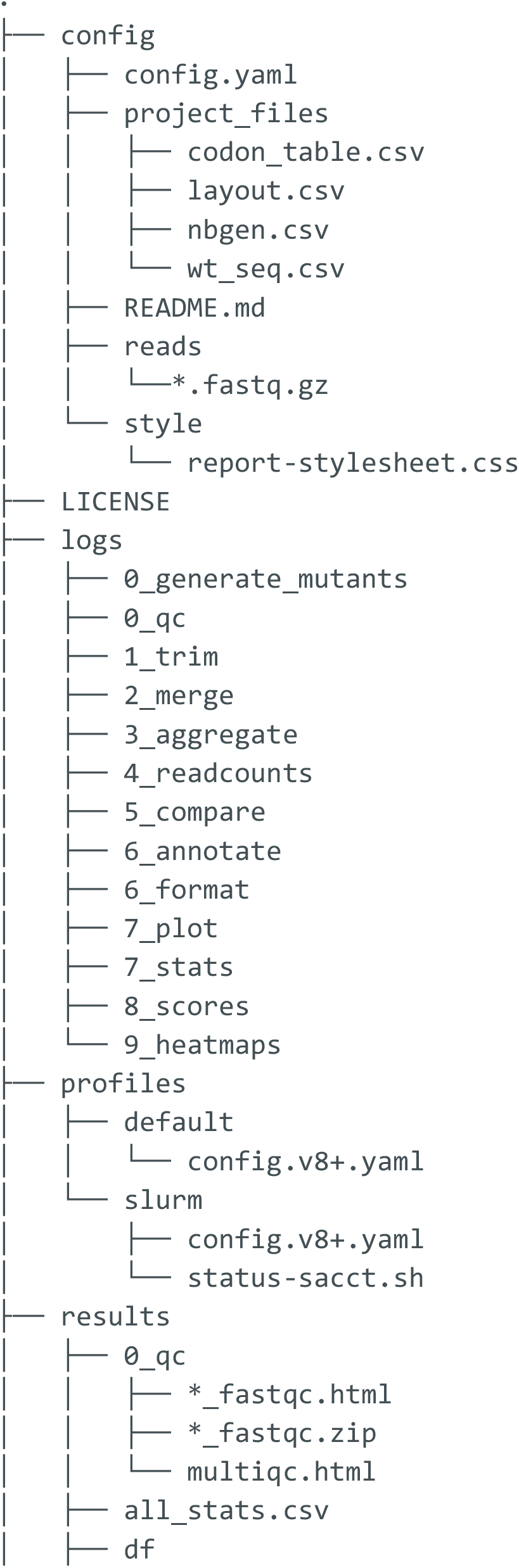
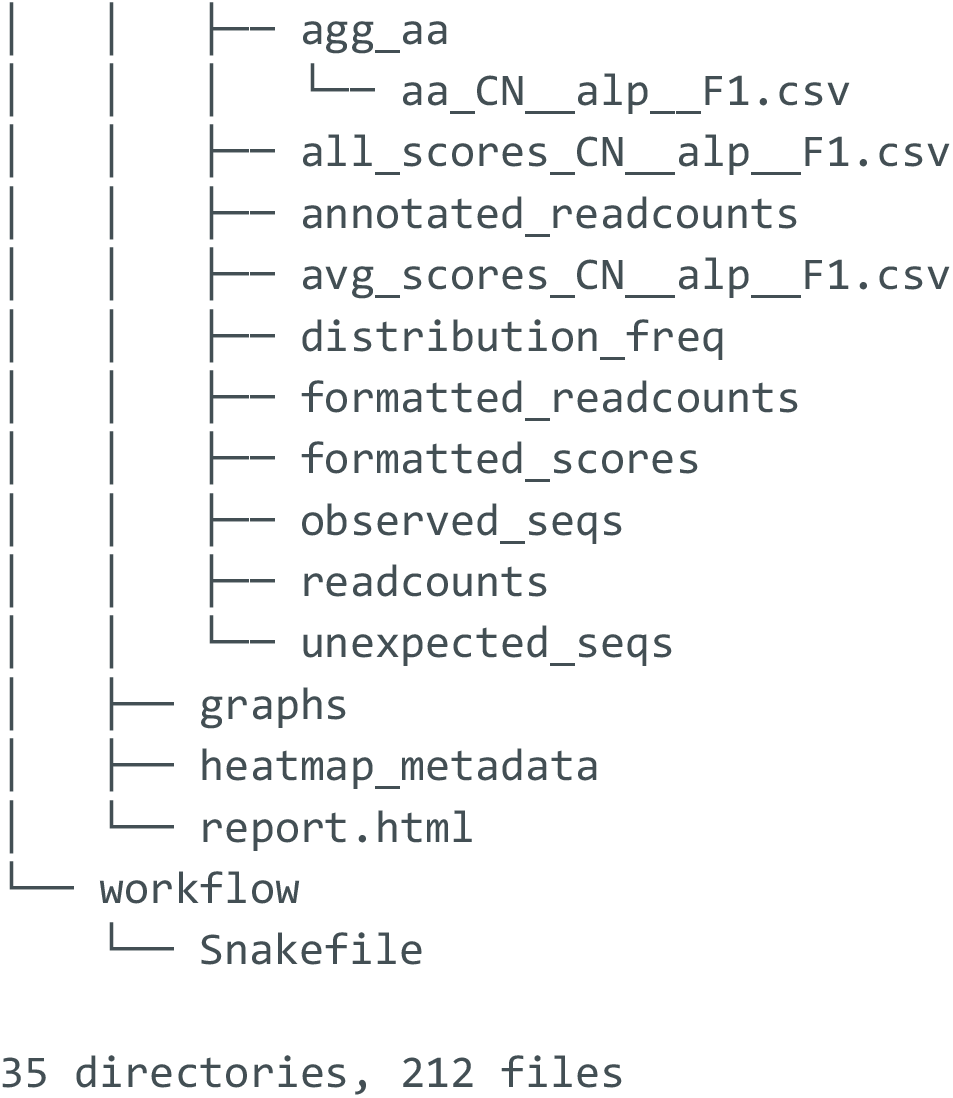
File tree upon full completion of the workflow. This specific example was obtained with the toy dataset. Some directories are collapsed for clarity. Some files (mostly FASTA files) are temporarily created for processing purposes but deleted as soon as the output from all dependent downstream rules has been generated. These files and the corresponding directories they are located in do not feature in this tree.

**Supplementary Figure 3.**
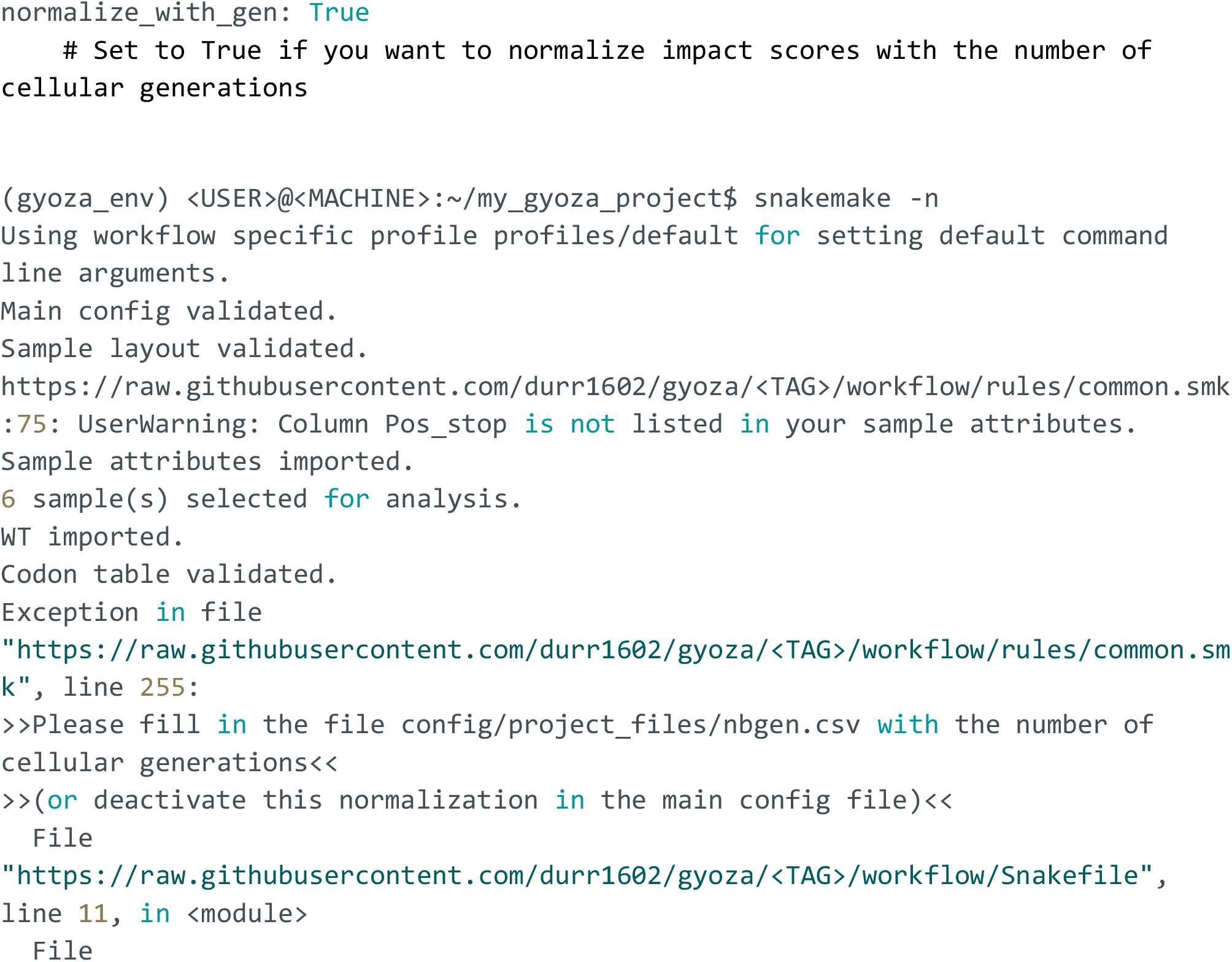
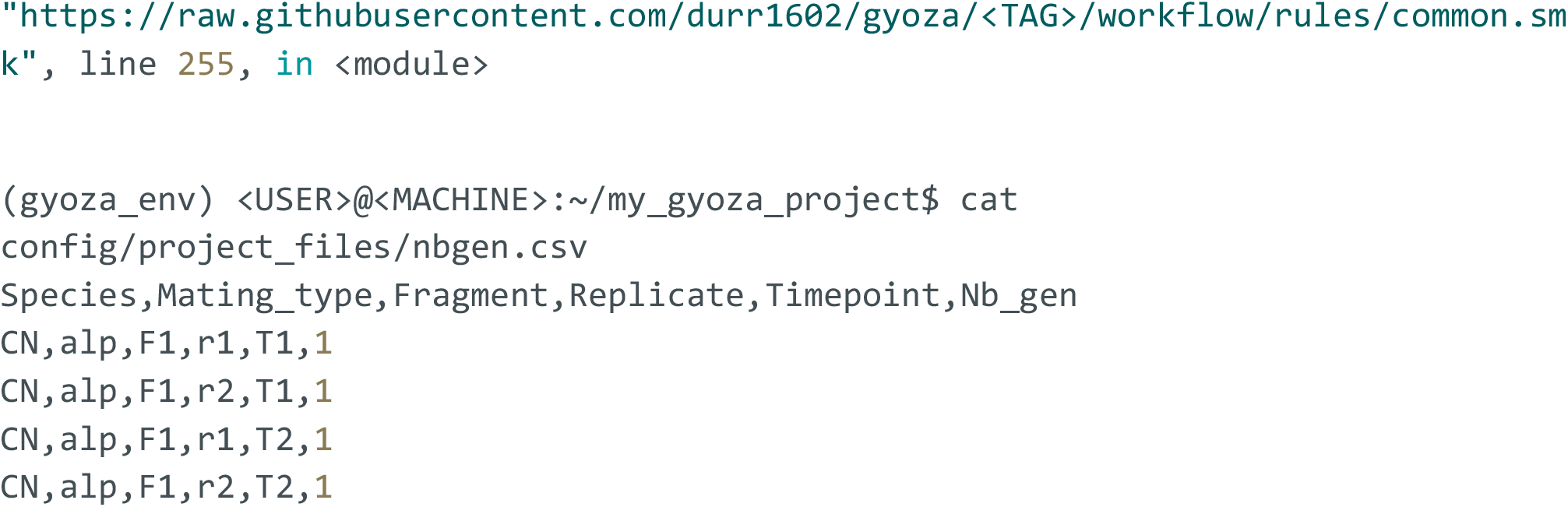
Normalization with the number of cellular generations. When opting in for normalization with the number of cellular generations in the config (first code block), the workflow generates a template file and prompts the user to fill it with numbers for each indicated condition (second block). In the corresponding file (third code block), we can see each combination of sample attributes (as defined in the config), replicate and time point arranged on different rows. For each row, the number of cellular generations between T0 and the indicated time point (on the same row) has been initialized to 1. From there, the user can either modify the file directly or use a custom script to generate one, making sure the format respects the template’s (it is not required to respect the same order of rows or columns). Normalization is disabled by default for the toy dataset. In order to specifically trigger this, the config was modified as shown in the first block and a dry run was re-executed after the initial one shown in Figure S1.

### Conda

At the time of writing, Conda is no longer a required dependency of Snakemake and can be replaced with Pixi. However, integrated package management still (again, currently) requires Conda. The main drawback is that it often comes with the installation of a large volume of files and may, in certain rare instances, interfere with already installed dependencies. It is one of the reasons why Conda is prohibited on some systems. Unfortunately, there is no easy fix for this. Until Snakemake allows the specification of predefined virtual environments, we believe the current solution offers the most interesting tradeoff between robustness and flexibility. Briefly, rule definitions must include a “conda” statement which points to a YAML requirements file (either locally in the “envs” directory, or online if the rule uses a wrapper). Each file lists the necessary packages with versions to be installed in a virtual Conda environment to ensure that there are no software incompatibilities. The first time the workflow is run, Snakemake will download and install all necessary packages in virtual Conda environments. This step might take time but is skipped if the environments already exist (e.g. if the workflow is run a second time). During the execution of the workflow, rule-dedicated environments are automatically activated.

